# PCH-2 collaborates with CMT-1 to proofread meiotic homolog interactions

**DOI:** 10.1101/780254

**Authors:** Stefani Giacopazzi, Daniel Vong, Alice Devigne, Needhi Bhalla

**Author notes:** these authors contributed equally (Stefani Giacopazzi, Daniel Vong). corresponding author, Department of Molecular, Cell and Developmental Biology, 225 Sinsheimer Labs, University of California, Santa Cruz, Santa Cruz, CA 95064.

## Abstract

The conserved ATPase, PCH-2/TRIP13, is required during both the spindle checkpoint and meiotic prophase. However, it’s specific role in regulating meiotic homolog pairing, synapsis and recombination has been enigmatic. Here, we report that this enzyme is required to proofread meiotic homolog interactions. We generated a mutant version of PCH-2 in *C. elegans* that binds ATP but cannot hydrolyze it: *pch-2*^*E253Q*^. *In vitro*, this mutant binds its substrates but is unable to remodel them. This mutation results in non-homologous synapsis and loss of crossover assurance. Surprisingly, worms with a null mutation in PCH-2’s adapter protein, CMT-1, the ortholog of p31^comet^, localize PCH-2 to meiotic chromosomes, exhibit non-homologous synapsis and lose crossover assurance. The similarity in phenotypes between *cmt-1* and *pch-2*^*E253Q*^ mutants indicate that PCH-2 can bind its meiotic substrates in the absence of CMT-1, in contrast to its role during the spindle checkpoint, but requires its adapter to hydrolyze ATP and remodel them.

## Introduction

Sexual reproduction relies on meiosis, the specialized cell division that generates haploid gametes, such as sperm and eggs, from diploid progenitors so that fertilization restores diploidy. To ensure that gametes inherit the correct number of chromosomes, meiotic chromosome segregation is exquisitely choreographed: Homologous chromosomes segregate in meiosis I and sister chromatids segregate in meiosis II. Having an incorrect number of chromosomes, also called aneuploidy, is associated with infertility, miscarriages and birth defects, underscoring the importance of understanding this process to human health.

Events in meiotic prophase ensure proper chromosome segregation. During prophase, homologous chromosomes undergo progressively intimate interactions that culminate in synapsis and crossover recombination (reviewed in [1]). After homologs pair, a macromolecular complex, called the synaptonemal complex (SC), is assembled between them in a process called synapsis. Synapsis is a prerequisite for crossover recombination to generate the linkages, or chiasmata, between homologous chromosomes that direct meiotic chromosome segregation. Defects in any of these events can result in chromosome missegregation during the meiotic divisions and gametes, and therefore embryos, with an incorrect number of chromosomes.

The conserved AAA-ATPase PCH-2 (Pch2 in budding yeast, PCH2 in Arabidopsis and TRIP13 in mice) is crucial to coordinate these events in meiotic prophase. *In vitro* and cytological experiments indicate that it does this by using the energy of ATP hydrolysis to remodel meiotic HORMADs [2–6], chromosomal proteins that are essential for pairing, synapsis, recombination and checkpoint function [7–18]. HORMADs are a protein family defined by the ability of a domain, the HORMA domain, to adopt multiple conformations that specify protein function [19–21]. Meiotic HORMADs appear to adopt two conformations: a “closed” version, that it forms upon binding a short peptide, called the closure motif, within other proteins [22]; and a more extended, or “unlocked,” version [23]. The closed form of meiotic HORMADs assemble on meiotic chromosomes to drive pairing, synapsis and recombination [22]. The physiological relevance of the unlocked version is unknown.

Cytological experiments in budding yeast, plants and mice show that PCH-2 removes or redistributes meiotic HORMADs on chromosomes [2, 4–6]. This change in localization both contributes to and reflects the progression of meiotic prophase events [2, 4–6]. However, major events in meiotic prophase, such as pairing, synapsis initiation and recombination, precede the removal or redistribution of meiotic HORMADs, raising the question of how PCH-2 affects these events to produce the phenotypes reported in its absence. Indeed, the phenotypes associated with loss of Pch2 are inconsistent with its only role being the redistribution of meiotic HORMADs [24–26]. In *C. elegans*, PCH-2 regulates meiotic prophase events but not the localization of meiotic HORMADs [27], indicating that 1) the removal or relocalization of meiotic HORMADs is not essential for PCH-2’s effects on pairing, synapsis and recombination; and 2) this model organism is particularly relevant to better understand how PCH-2 regulates these events.

We previously showed that in the absence of PCH-2, homolog pairing, synapsis and recombination are accelerated and associated with an increase in meiotic defects, leading us to speculate that PCH-2 disassembles molecular intermediates that underlie these events to ensure their fidelity [27]. Specifically, we hypothesized that these molecular intermediates involve meiotic HORMADs and PCH-2 remodels meiotic HORMADs to proofread these homolog interactions [27]. To test this model, we generated a mutant version of PCH-2 that binds ATP, and its substrates, but cannot hydrolyze ATP and therefore cannot remodel and release its substrates. We reasoned that if our hypothesis is correct, this mutant protein will “trap,” and allow us to more easily visualize, inappropriate homolog interactions that are incorrectly stabilized. This is accurate: *pch-2*^*E253Q*^ mutants delay homolog pairing and accelerate synapsis, producing non-homologous synapsis. We see a similar effect on recombination. Meiotic DNA repair occurs with normal kinetics but crossover assurance is lost, suggesting that crossover-specific recombination intermediates between incorrect partners may also be inappropriately stabilized. Surprisingly, loss of CMT-1, an adapter protein that is thought to be essential for PCH-2 to bind its substrates, phenotypically resembles *pch-2*^*E253Q*^ mutants. Since PCH-2 localizes normally to meiotic chromosomes in the absence of CMT-1, these data indicate that CMT-1 is dispensable for PCH-2 to bind its meiotic substrates on chromosomes but is essential to hydrolyze ATP and remodel its substrates.

## Results

### PCH-2^E253Q^ localizes normally during meiotic prophase

We introduced a mutation by CRISPR/Cas9 genomic editing [28, 29] in the Walker B motif of PCH-2, E253Q, that allows it to bind ATP, but not hydrolyze it. We designated this allele *pch-2(blt5)* but will refer to it as *pch-2*^*E253Q*^. In budding yeast, this mutation abolishes PCH-2’s *in vivo* meiotic function [3]. *In vitro*, PCH-2^E253Q^ retains high affinity nucleotide binding, forms stable hexamers and interacts with its substrates [30]. In meiotic nuclei, PCH-2^E253Q^ localized normally to meiotic chromosomes (Figure 1A), appearing as foci just prior to the entry into meiosis (also known as the transition zone) and co-localizing with the synaptonemal complex once synapsis had initiated (Figure 1B). It was also removed from the synaptonemal complex in late pachytene, similar to wildtype PCH-2 (Figure 1C).

**Figure 1:**
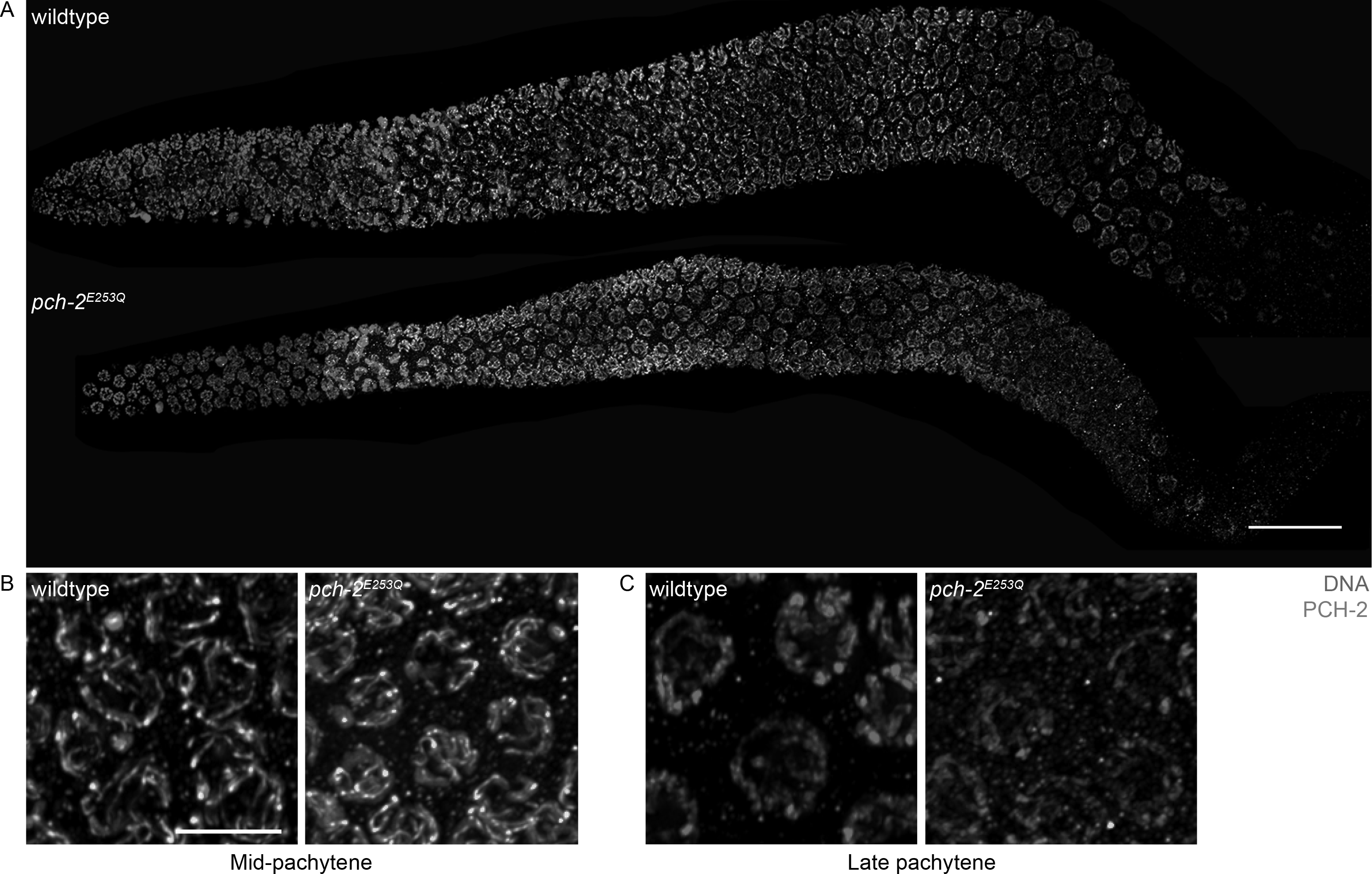
PCH-2^E253Q^ localizes to meiotic chromosomes similar to wildtype PCH-2. A. Whole germline images of PCH-2 and DAPI staining in a wildtype and *pch-2*^*E253Q*^ mutant germline. Scale bar indicates 20 microns. Meiotic nuclei in mid-pachytene (B) and late pachytene (C) stained with DAPI and antibodies against PCH-2 in wildtype animals and *pch-2*^*E253Q*^ mutants. Unless otherwise stated, all scale bars indicate 5 microns.

### *pch-2*^*E253Q*^ mutants delay pairing

We analyzed pairing in *syp-1* mutants. SYP-1 is a component of the synaptonemal complex (SC) and *syp-1* mutants fail to assemble SC between paired homologs, allowing us to more easily visualize pairing intermediates in the absence of synapsis [31]. In *C. elegans*, cis-acting sites called pairing centers (PCs) are required for pairing and synapsis [32]. In the absence of synapsis, homologous chromosomes exhibit stable pairing of PC, but not non-PC, ends of chromosomes [32]. We monitor synapsis-independent pairing at PC sites by performing immunofluorescence against HIM-8, which binds PCs of X chromosomes [33] (Figure 2B). A single focus of HIM-8 indicates paired X chromosomes while two foci indicate unpaired X chromosomes. When we image stained germlines, we divide germlines into six equal sized zones (see cartoon in Figure 2A). Because meiotic nuclei are arranged spatiotemporally in the germline, this allows us to assess pairing as a function of meiotic progression.

**Figure 2:**
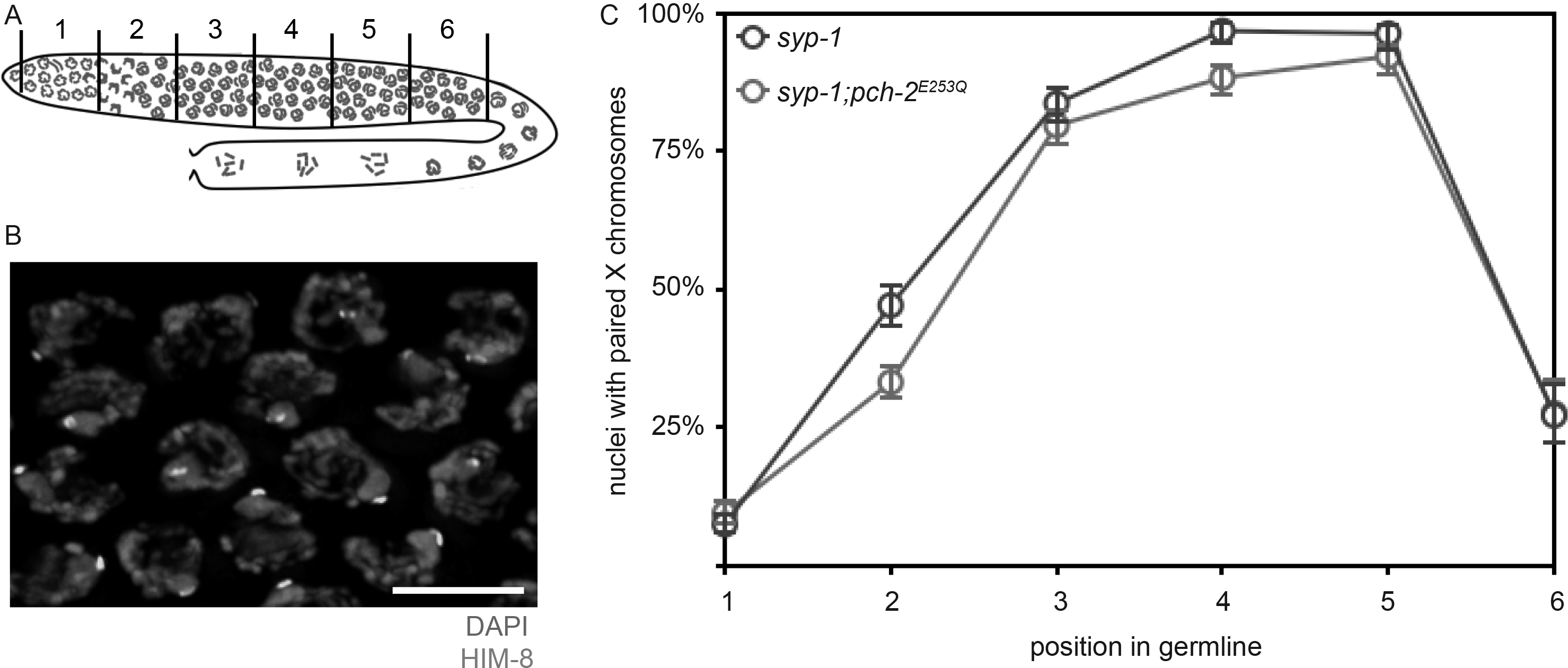
Homolog pairing is delayed in *pch-2*^*E253Q*^ mutants. A. Cartoon of *C. elegans* germline divided into six equivalent zones. B. Images of meiotic nuclei stained with DAPI and antibodies against HIM-8. C. Timecourse of X chromosome pairing in wildtype and *pch-2*^*E253Q*^ mutant germlines.

*syp-1* mutants initiate pairing in zone 2, maintain pairing at the X chromosome PC in zones 3,4 and 5 and destabilize pairing in zone 6 (Figure 2C). *pch-2*^*E253Q*^*;syp-1* double mutants also initiate PC pairing in zone 2 but have significantly fewer nuclei with paired HIM-8 signals than *syp-1* single mutants. In later zones, *pch-2*^*E253Q*^*;syp-1* mutants more closely resemble *syp-1* single mutants. Thus *pch-2*^*E253Q*^*;syp-1* mutants delay pairing at PCs, consistent with our model that PCH-2 proofreads homolog pairing intermediates: If *pch-2*^*E253Q*^ mutants “trap” incorrect pairing intermediates, between non-homologous chromosomes, for example, this would delay accurate pairing between homologous chromosomes.

### *pch-2*^*E253Q*^ mutants accelerate synapsis and produce non-homologous synapsis

Next, we assayed synapsis by monitoring the colocalization of two SC components, HTP-3 and SYP-1 [31, 32]. When HTP-3 and SYP-1 colocalize, chromosomes are synapsed while regions of HTP-3 devoid of SYP-1 are unsynapsed (see arrows in Figure 3A). We evaluated the percentage of nuclei that had completed synapsis as a function of meiotic progression (Figure 3B). In contrast, to our analysis of pairing, synapsis in *pch-2*^*E253Q*^ mutants was accelerated, similar to *pch-2Δ* mutants [27]: More nuclei had completely synapsed chromosome when synapsis initiates in zone 3 in *pch-2*^*E253Q*^ mutants than control germlines (Figure 3B). In contrast to *pch-2Δ* mutants [27], *pch-2*^*E253Q*^ mutants did not exhibit defects in SC disassembly in zone 6 (Figure 3B).

**Figure 3:**
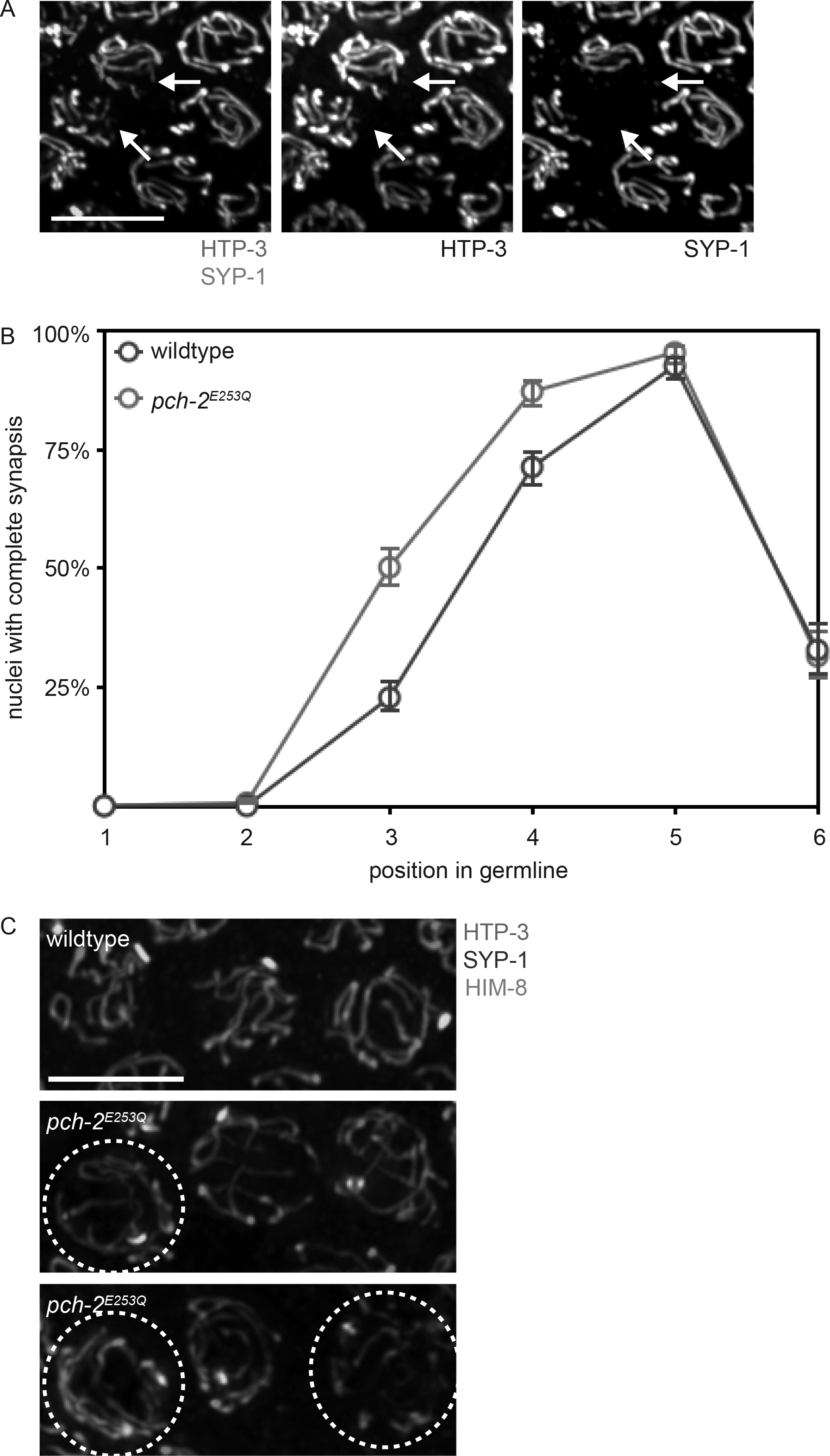
Synapsis is accelerated in *pch-2*^*E253Q*^ mutants, producing non-homologous synapsis. A. Images of meiotic nuclei stained with antibodies against HTP-3 and SYP-1. Color and grey scale images are provided. Arrows indicate unsynapsed chromosomes. B. Timecourse of synapsis in wildtype and *pch-2*^*E253Q*^ mutant germlines. C. Images of meiotic nuclei stained with antibodies against HTP-3, SYP-1 and HIM-8 in wildtype animals and *pch-2*^*E253Q*^ mutants. Circled nuclei have undergone non-homologous synapsis.

We reasoned that the delay in pairing, combined with the acceleration in synapsis, raised the possibility that these events, which are typically coupled, had been uncoupled. To test this possibility, we evaluated whether non-homologous synapsis occurs in *pch-2*^*E253Q*^ mutants. We stained *pch-2*^*E253Q*^ mutants with antibodies against SC components HTP-3 and SYP-1 to evaluate synapsis and HIM-8 to assess pairing. All nuclei in wildtype animals and most nuclei in *pch-2*^*E253Q*^ mutants exhibited homologous synapsis (uncircled nuclei in Figure 3C). However, we identified meiotic nuclei in which all chromosomes were synapsed but contained two HIM-8 foci, indicating these chromosomes had synapsed with non-homologous partners (circled nuclei in Figure 3C). We also observed non-homologous synapsis when we monitored pairing of autosomes using antibodies against ZIM-1 and ZIM-2, proteins that bind autosomal PCs [34] (data not shown). When we quantified this defect, we detected non-homologous synapsis of X chromosomes in 3% of meiotic nuclei (data not shown). Since *C. elegans* have six pairs of homologous chromosomes, it’s possible that as many as 18% of meiotic nuclei have at least one pair of chromosomes undergoing non-homologous synapsis in *pch-2*^*E253Q*^ mutants.

### *pch-2*^*E253Q*^ mutants lose crossover assurance

We then monitored meiotic DNA repair and the formation of crossovers. To assess DNA repair, we monitor the appearance and disappearance of RAD-51 from meiotic chromosomes as a function of meiotic progression (Figure 4A). RAD-51 is required for all meiotic DNA repair in *C. elegans* [35]. Its appearance on chromosomes signals the formation of double strand breaks (DSBs) that need to be repaired (see zones 3 and 4 in wildtype in Figure 4B) and its disappearance indicates the entry of DSBs into a DNA repair pathway (see zones 5 and 6 in wildtype in Figure 4B) [36]. The appearance and disappearance of RAD-51 on meiotic chromosomes in *pch-2*^*E253Q*^ mutants was indistinguishable from that in wildtype in zones 1-4. In zone 5, we detected fewer RAD-51 foci, suggesting that meiotic DNA repair may occur slightly more rapidly in *pch-2*^*E253Q*^ mutants. However, this acceleration is less severe than what we observe in *pch-2Δ* mutants [27].

**Figure 4:**
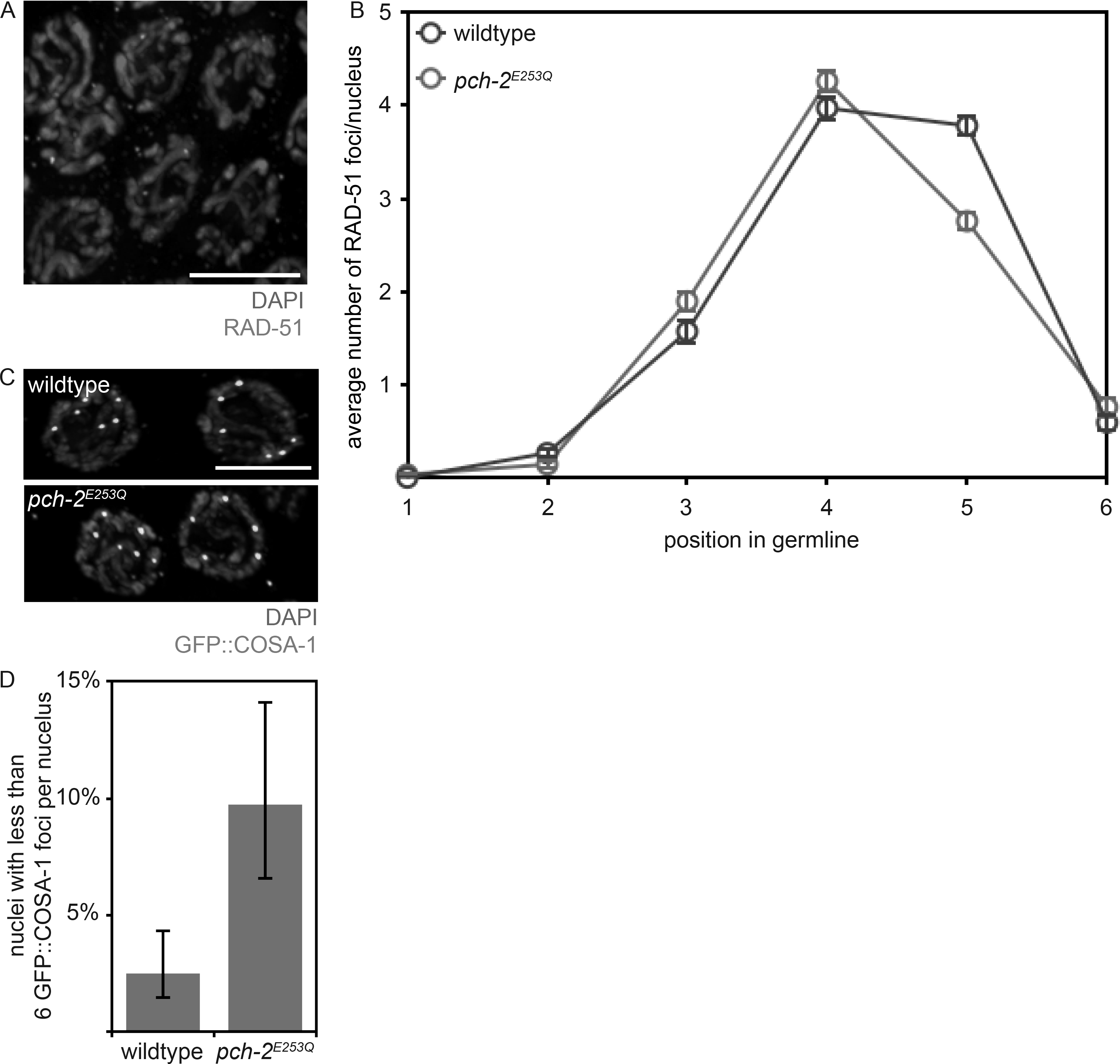
Meiotic DNA repair is not affected in *pch-2*^*E253Q*^ mutants but crossover assurance is. A. Images of meiotic nuclei stained with DAPI and antibodies against RAD-51. B. Timecourse of the average number of RAD-51 foci per nucleus in wildtype and *pch-2*^*E253Q*^ mutant germlines. C. Images of meiotic nuclei stained with DAPI and antibodies against GFP. D. Percentage of nuclei with less than six GFP∷COSA-1 foci in wildtype animals and *pch-2*^*E253Q*^mutants.

We assayed crossover formation in *pch-2*^*E253Q*^ mutants by visualizing GFP∷COSA-1 foci formation in late meiotic prophase. GFP∷COSA-1 cytologically marks putative crossovers and its appearance as robust foci is mechanistically associated with the process of crossover designation [37]. Almost all nuclei in wildtype worms have six GFP∷COSA-1 foci per nucleus, one per homolog pair (Figure 4C, top, and 4D) [37]. In contrast, *pch-2*^*E253Q*^ mutants have a greater number of nuclei with less than six GFP∷COSA-1 foci, indicating a defect in crossover assurance (Figure 4C, bottom, and 4D). Given the *in vitro* behavior of the PCH-2^E253Q^ hexamer [30], this phenotype is likely because inappropriate crossover intermediates are stabilized at the expense of appropriate ones. Alternatively, these meiotic nuclei may result from the non-homologous synapsis we also observe in this mutant background. However, we do not observe a delay in meiotic DNA repair in *pch-2*^*E253Q*^ mutants (Figure 4B). Further, *pch-2*^*E253Q*^ mutants do not activate feedback mechanisms, as visualized by the extension of DSB-1 on meiotic chromosomes [38, 39] (Figure S1). Therefore, we favor the first interpretation.

### CMT-1 is required for the synapsis checkpoint

During the spindle checkpoint, PCH-2 requires the presence of an adapter protein, CMT-1 (p31^comet^ in mammalian cells), to bind and remodel a HORMAD protein required for the spindle checkpoint response, Mad2 [30, 40, 41]. Based on a recent report that the rice ortholog of CMT-1 is an SC component and required for pairing, synapsis and double strand break formation [42], we tested whether *cmt-1* had a role in meiotic prophase by assessing whether it was required for meiotic prophase checkpoints in *C. elegans*. We introduced a null mutation of *cmt-1* into *syp-1* mutants. The absence of SC in *syp-1* mutants results in unsynapsed chromosomes, which activate very high levels of germline apoptosis in response to both the synapsis and the DNA damage checkpoint [43] (Figure 5A). *cmt-1* single mutants have wildtype levels of apoptosis (Figure 5B). In contrast to *syp-1* single mutants, *syp-1;cmt-1* double mutants had intermediate levels of germline apoptosis (Figure 5B), indicating that CMT-1 is required for either the synapsis or DNA damage checkpoint. To distinguish between these two checkpoints, we use *spo-11; syp-1* double mutants, which do not generate double strand breaks [44]. As a result, these double mutants do not activate the DNA damage checkpoint (Figure 5A) and produce an intermediate level of apoptosis (Figure 5B). [43]. When we generate *cmt-1; spo-11; syp-1* triple mutants, we observe wildtype levels of apoptosis, similar to *cmt-1* single mutants (Figure 5B). Therefore, CMT-1 is required to activate apoptosis in response to the synapsis checkpoint, similar to *pch-2Δ* mutants.

**Figure 5:**
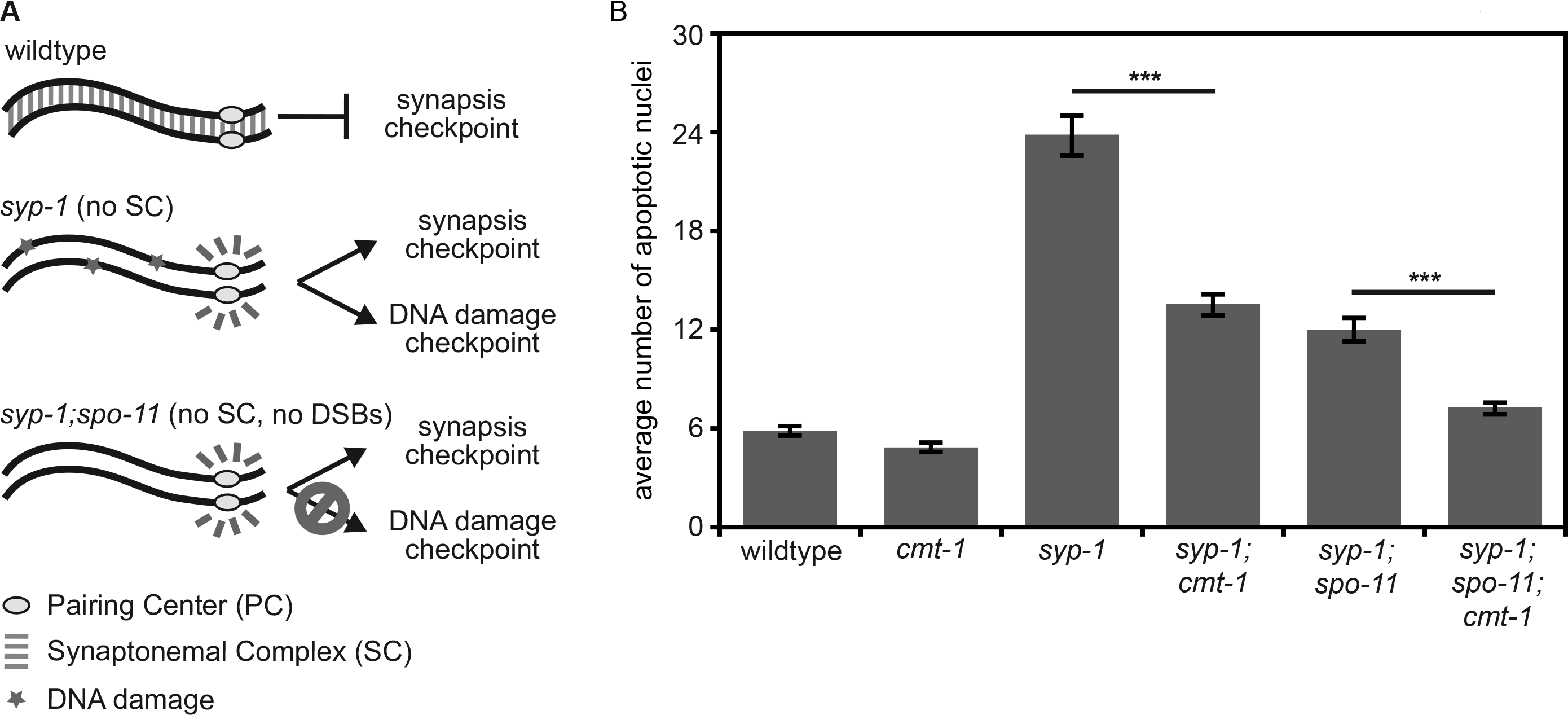
CMT-1 is required for the synapsis checkpoint. A. Cartoon of meiotic checkpoint activation in *C. elegans*. B. Mutation of *cmt-1* reduces apoptosis in *syp-1* single mutants and *syp-1;spo-11* double mutants. Error bars indicate 2XSEM. Significance was assessed by performing t-tests. A *** indicates a p value < 0.0001.

### *cmt-1* mutants phenotypically resemble *pch-2*^*E253Q*^ mutants

Based on the requirement for CMT-1 in the synapsis checkpoint, we assessed pairing, synapsis, meiotic DNA repair and crossover formation in *cmt-1* mutants. Similar to our analysis of *pch-2*^*E253Q*^ mutants, pairing was delayed in *cmt-1;syp-1* double mutants (Figure 6A) and synapsis was accelerated in *cmt-1* single mutants (Figure 6B). We also detected non-homologous synapsis at levels similar to that of *pch-2*^*E253Q*^ mutants (Figure 6C). Meiotic DNA repair in *cmt-1* mutants, as visualized by the appearance and disappearance of RAD-51 (Figure S2A), and meiotic progression, as visualized by the appearance and disappearance of a protein required for DSB formation, DSB-1 (Figure S1), closely resembled that of wildtype germlines. In addition, crossover assurance was reduced (Figure S2B). These data indicate that, unlike with Mad2, PCH-2 can bind its meiotic substrates effectively in the absence of CMT-1 but CMT-1 is required for PCH-2’s hydrolysis of ATP. Consistent with this interpretation, *pch-2*^*E253Q*^*;cmt-1* double mutants have a similar frequency of non-homologous synapsis and nuclei with less than six GFP∷COSA-1 foci as either single mutant (data not shown). Further, since *cmt-1* mutants result in meiotic phenotypes distinct from those we reported for spindle checkpoint mutants [45], specifically non-homologous synapsis, CMT-1’s role in meiotic prophase is independent of its regulation of Mad2.

**Figure 6:**
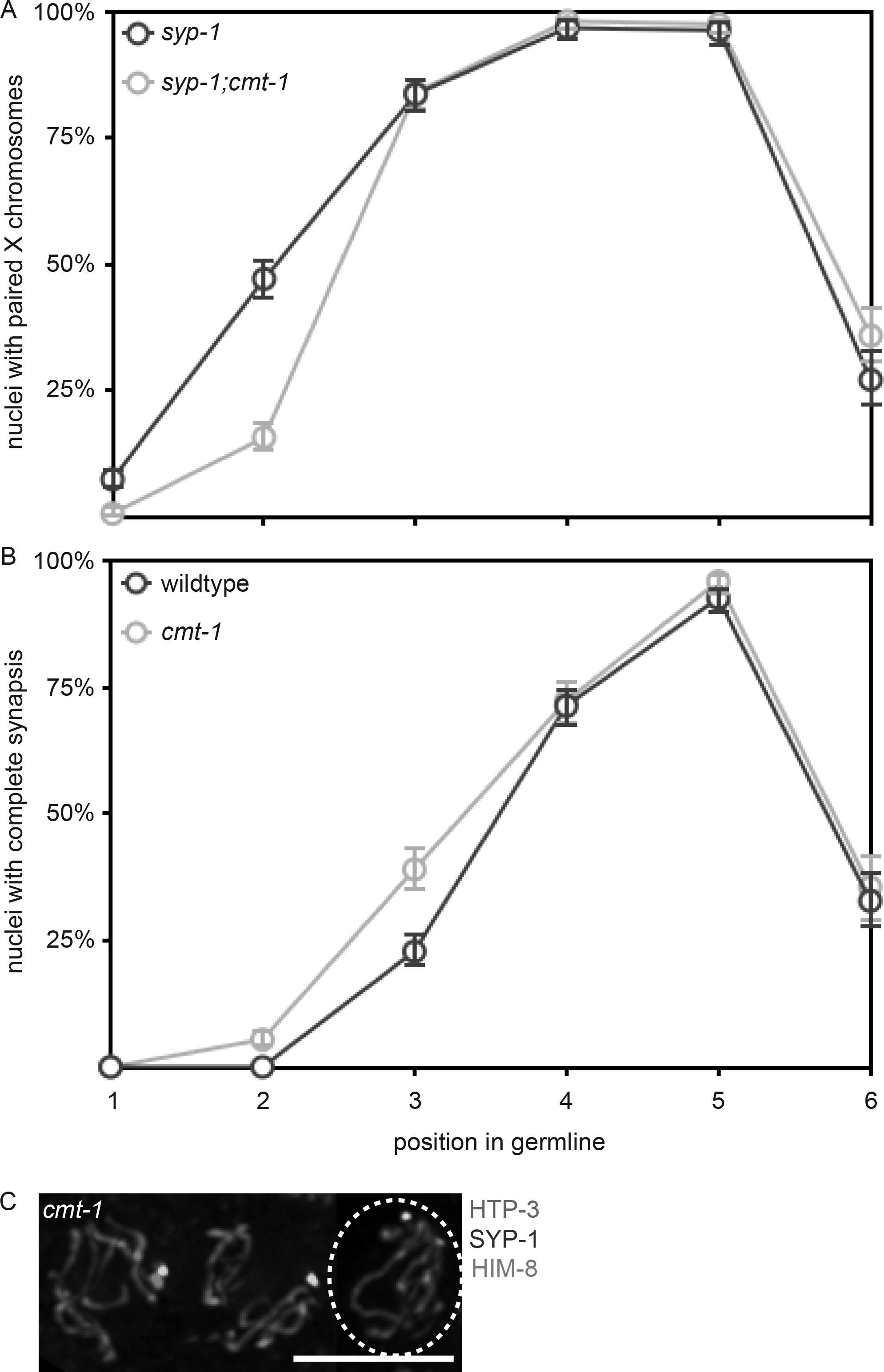
*cmt-1* mutants delay pairing, accelerate synapsis and exhibit non-homologous synapsis, similar to *pch-2*^*E253Q*^ mutants. A. Timecourse of pairing in wildtype and *cmt-1* mutant germlines. B. Timecourse of synapsis in wildtype and *cmt-1* mutant germlines. C. Images of meiotic nuclei stained with antibodies against HTP-3, SYP-1 and HIM-8 in *cmt-1* mutants. The circled nucleus has undergone non-homologous synapsis.

### Non-homologous synapsis in *cmt-1* mutants relies on PCH-2

We previously showed that CMT-1 is required for PCH-2 to localize to mitotic chromosomes during the spindle checkpoint [46]. We tested whether this was also true during meiotic prophase and found that CMT-1 was dispensable for PCH-2’s localization to meiotic chromosomes (Figure 7A). Consistent with CMT-1 being required for PCH-2’s hydrolysis of ATP and release of substrates, we observed that PCH-2 stayed on meiotic chromosomes slightly longer than in wildtype germlines (Figure 7C).

**Figure 7:**
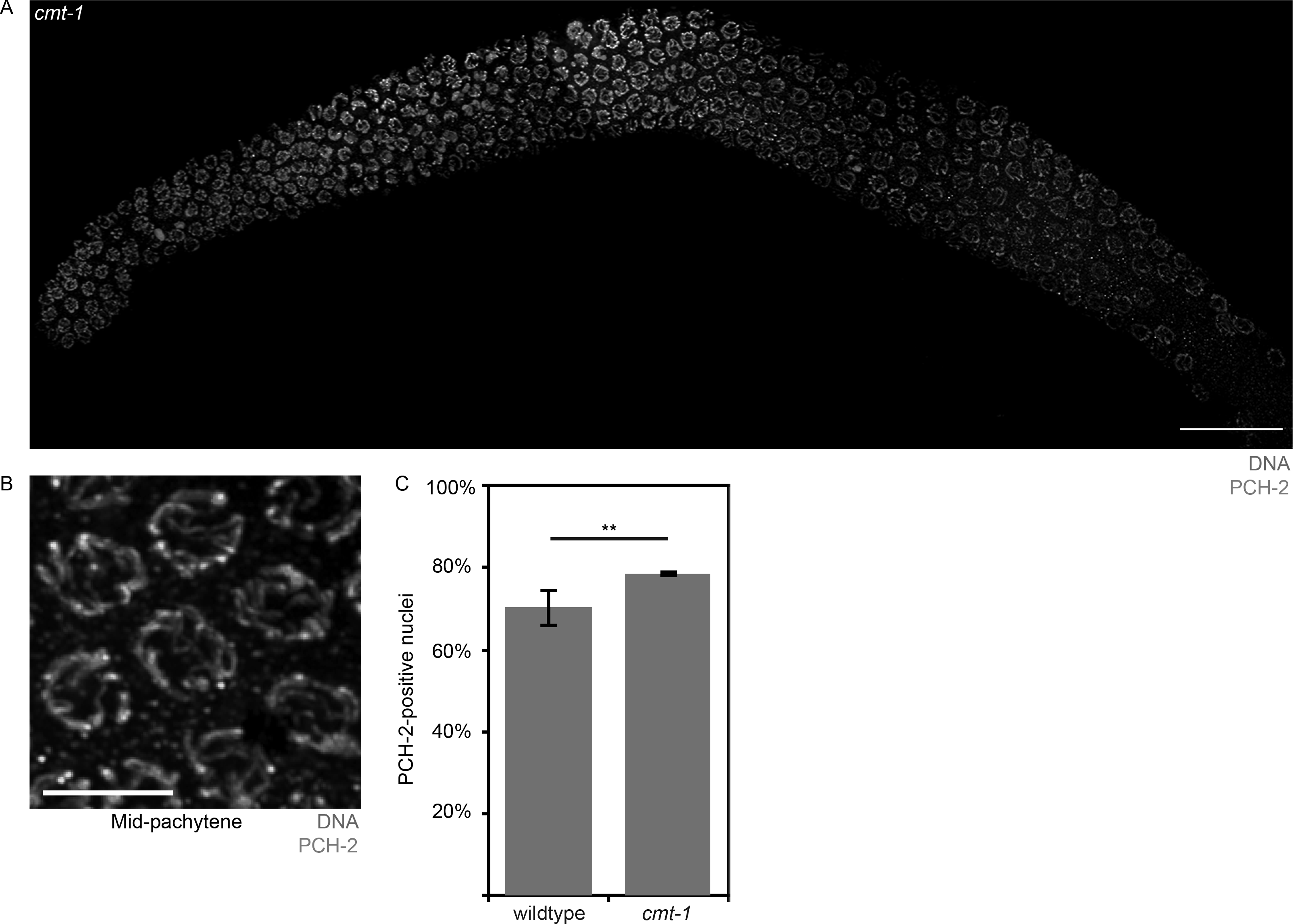
PCH-2 localizes to meiotic chromosomes in *cmt-1* mutants. A. Whole germline image of PCH-2 and DAPI staining in a *cmt-1* mutant germline. Scale bar indicates 20 microns. B. Meiotic nuclei in mid-pachytene stained with DAPI and antibodies against PCH-2 in *cmt-1* mutants. C. Quantification of percentage of PCH-2-positive nuclei in wildtype and *cmt-1* mutant germlines. Significance was assessed by performing t-tests. A ** indicates a p value < 0.01.

During mitotic divisions in the *C. elegans* embryo, loss of CMT-1 partially suppresses the spindle checkpoint defect observed in *pch-2* mutants [46]. Since PCH-2 ensures availability of the inactive conformer of Mad2 so that it can be converted to the active one during checkpoint activation [47, 48], we hypothesize that this genetic interaction results because loss of CMT-1 makes more inactive Mad2 available. In this way, PCH-2 and CMT-1 would control the spindle checkpoint by regulating available, soluble pools of inactive and active Mad2. We wondered if there was a similar relationship between PCH-2 and CMT-1 in meiotic prophase. To test this, we generated *pch-2Δ;cmt-1* double mutants and assayed non-homologous synapsis. Unlike *cmt-1* single mutants and similar to *pch-2Δ* mutants, we did not detect non-homologous synapsis in *pch-2Δ;cmt-1* double mutants. Thus, PCH-2 is epistatic to CMT-1 during meiosis, at least in the context of non-homologous synapsis, supporting our interpretation that PCH-2 can bind its meiotic substrates independent of CMT-1 and suggesting that some of PCH-2’s regulation of its meiotic substrates occurs directly on chromosomes.

## Discussion

*In vitro*, PCH-2^E253Q^ protein binds ATP and its spindle checkpoint substrate, Mad2, but fails to remodel it, due to an inability to hydrolyze ATP [30]. Here, we show that the meiotic phenotypes observed in *pch-2*^*E253Q*^ mutants are consistent with a role for PCH-2 in disassembling inappropriate pairing, synapsis and crossover recombination intermediates in *C. elegans*, resulting in non-homologous synapsis (Figure 3C) and a loss of crossover assurance (Figure 4D). The effect we observe on these meiotic prophase events seems limited, affecting a small proportion of nuclei, but this may reflect either the inherent fidelity of these processes or the existence of redundant mechanisms that contribute to fidelity. Further, a role for PCH-2 in proofreading may be more apparent in situations in which homolog interactions are more challenging. Indeed, PCH-2’s involvement in the interchromosomal effect [49], in which chromosome rearrangements affect crossover control on other chromosomes, supports this possibility. Nevertheless, our results demonstrate that PCH-2 proofreads interactions between homologous chromosomes to ensure that they are correct.

We were surprised to see that *cmt-1* mutants localized PCH-2 to meiotic chromosomes (Figure 7), unlike what we observe during the spindle checkpoint response [46], and closely resembled *pch-2*^*E253Q*^ mutants in exhibiting non-homologous synapsis and a loss of crossover assurance (Figures 6 and S2). These data indicate that PCH-2 is competent to binds its proposed meiotic substrates, meiotic HORMADs, in the absence of CMT-1 but that CMT-1 promotes PCH-2’s ATPase activity and ability to remodel these substrates. This is unlike its role in the spindle checkpoint, in which CMT-1/p31^comet^ is essential for PCH-2 to bind its substrate, Mad2 [30, 40, 41, 50]. This difference raises the possibility that the interaction between PCH-2 and meiotic HORMADs during their remodeling is substantially different than that with Mad2. This hypothesis is supported by the findings that budding yeast, in which Pch2 was originally identified [51], does not have a CMT-1/p31^comet^ ortholog [52, 53] and budding yeast Pch2 can directly interact with and remodel the budding yeast meiotic HORMAD, Hop1, *in vitro* [3].

The PCH-2/HORMAD genetic module is evolutionarily ancient, having been identified as operons in several bacteria [54], suggesting that this pair of proteins has been co-opted to function in multiple molecular contexts. While we can easily detect an interaction between PCH-2 and CMT-1, and CMT-1 and Mad2, by two-hybrid experiments [46], we have not been able to observe a similar interaction between PCH-2 or CMT-1 and any of the four meiotic HORMADs present in *C. elegans*, HTP-3, HIM-3, HTP-1 and HTP-2 (data not shown), making it difficult to identify what regions of these proteins are required to interact with PCH-2. Genetic mutations, combined with cytological analysis, seems the most straightforward way, at least in *C. elegans*, to understand whether and how PCH-2 and meiotic HORMADs interact to regulate meiotic pairing, synapsis and recombination.

## Materials and Methods

### Genetics and worm strains

The *C. elegans* Bristol N2 [55] was used as the wild-type strain. All strains were maintained at 20°C under standard conditions unless otherwise stated. Mutations and rearrangements used were as follows:

LG I: *cmt-1(ok2879)*
LG II: *pch-2(blt5)*, *meIs8 [Ppie-1∷GFP∷cosa-1 + unc-119(+)]*
LG IV: *spo-11(ok79), nT1[unc-?(n754) let-?(m435)] (IV, V)*, *nTI [qIs51]*
LG V: *syp-1(me17)*, *bcIs39 [lim-7p∷ced-1∷GFP + lin-15(+)]*

The *pch-2(blt5)* allele, referred to as *pch-2*^*E253Q*^, was created by CRISPR-mediated genomic editing as described in [28, 29]. pDD162 was mutagenized using Q5 mutagenesis (New England Biolabs) and oligos GTTTTTGTTCTTATCGACGGTTTTAGAGCTAGAAATAGCAAGT and CAAGACATCTCGCAATAGG. The resulting plasmid was sequenced and two different correct clones (50ng/ul total) were mixed with pRF4 (100ng/ul) and the repair oligo CTCGTTCAAAAAATGTTCGATCAAATTGATGAACTAGCTGAAGATGAGAAGTGCATGGTTTTTGTGCTCATCGACCAAGTTTGATTTTTTTAAAAAACAATTTTTCTGGTTTTCATCAGTTTTTATGTCAGGTTGAAT (30ng/ul). Wildtype worms were picked as L4s, allowed to age 15-20 hours at 20°C and injected with the described mix. Worms that produced rolling progeny were identified and F1 rollers, as well as their wildtype siblings, were placed on plates seeded with OP50, 1-2 rollers per plate and 6-8 non-rolling siblings per plate, and allowed to produce progeny. PCR and Bsp1286I digestions were performed on these F1s to identify worms that contained the mutant allele and individual F2s were picked to identify mutant homozygotes. Multiple homozygotes carrying the *pch-2(blt5)* mutant allele were backcrossed against wildtype worms at least three times and analyzed to determine whether they produced the same mutant phenotype.

Scoring of germline apoptosis was performed as previously described in [43] with the following exceptions. L4 hermaphrodites were allowed to age for 22 hours. They were then mounted under coverslips on 1.5% agarose pads containing 0.2mM levamisole and scored. A minimum of twenty-five germlines was analyzed for each genotype.

### Antibodies, Immunostaining and Microscopy

DAPI staining and immunostaining was performed as in [43] 20 to 24 hours post L4 stage. Primary antibodies were as follows (dilutions are indicated in parentheses): rabbit anti-PCH-2 (1∶500) [27], rabbit anti-SYP-1 (1:500) [31], chicken anti-HTP-3 (1:250) [32], guinea pig anti-HTP-3 (1:250) [32], rabbit anti-ZIM-1 (1:1000), guinea pig anti-ZIM-2 (1:2500) [34], guinea pig anti-HIM-8 (1:250) [33], mouse anti-GFP (1:100) (Invitrogen), guinea pig anti-DSB-1 (1:500) [39] and rabbit anti-RAD-51 (1:1000) (Novus Biologicals). Secondary antibodies were Cy3 anti-rabbit, anti-guinea pig and anti-chicken (Jackson Immunochemicals) and Alexa-Fluor 488 anti-guinea pig and anti-rabbit (Invitrogen). All secondary antibodies were used at a dilution of 1:500.

All images were acquired using a DeltaVision Personal DV system (Applied Precision) equipped with a 100X N.A. 1.40 oil-immersion objective (Olympus), resulting in an effective XY pixel spacing of 0.064 or 0.040 μm. Three-dimensional image stacks were collected at 0.2-μm Z-spacing and processed by constrained, iterative deconvolution. Image scaling and analysis were performed using functions in the softWoRx software package. Projections were calculated by a maximum intensity algorithm. Composite images were assembled and some false coloring was performed with Adobe Photoshop.

Quantification of pairing, synapsis, RAD-51 foci, GFP∷COSA-1 foci, and DSB-1 positive nuclei was performed on animals 24 hours post L4 stage and with a minimum of three germlines per genotype. Relevant statistical analysis, as indicated in the Figure Legends, was used to assess significance.

---

**Figure S1:**
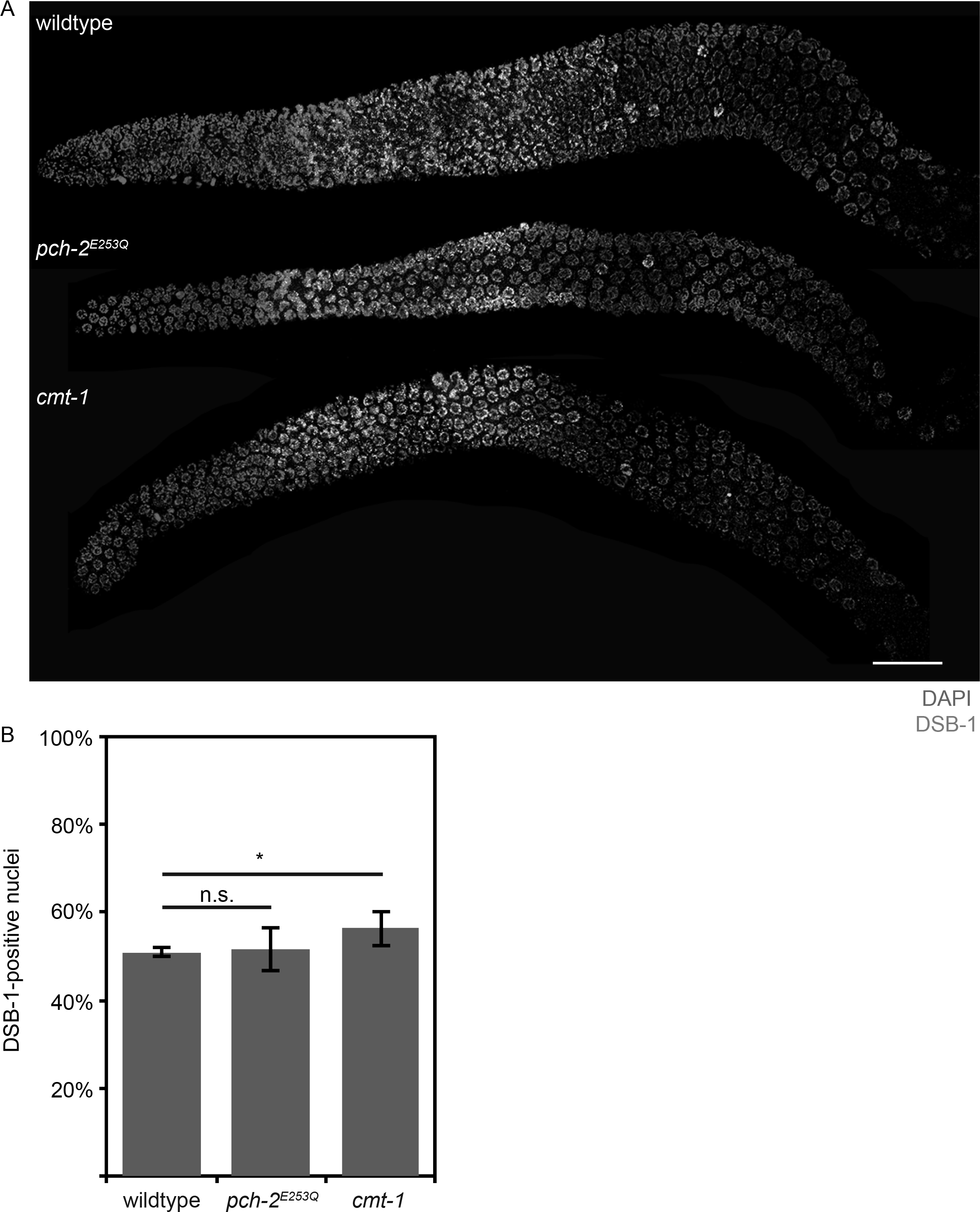
Meiotic progression is unaffected in *pch-2*^*E253Q*^ and *cmt-1* mutants. A. Whole germline images of DSB-1 and DAPI staining in a wildtype, *pch-2*^*E253Q*^ and *cmt-1* mutant germline. Scale bar indicates 20 microns. B. Quantification of percentage of DSB-1-positive nuclei in wildtype, *pch-2*^*E253Q*^ and *cmt-1* mutant germlines. Significance was assessed by performing t-tests. A * indicates a p value < 0.05.

**Figure S2:**
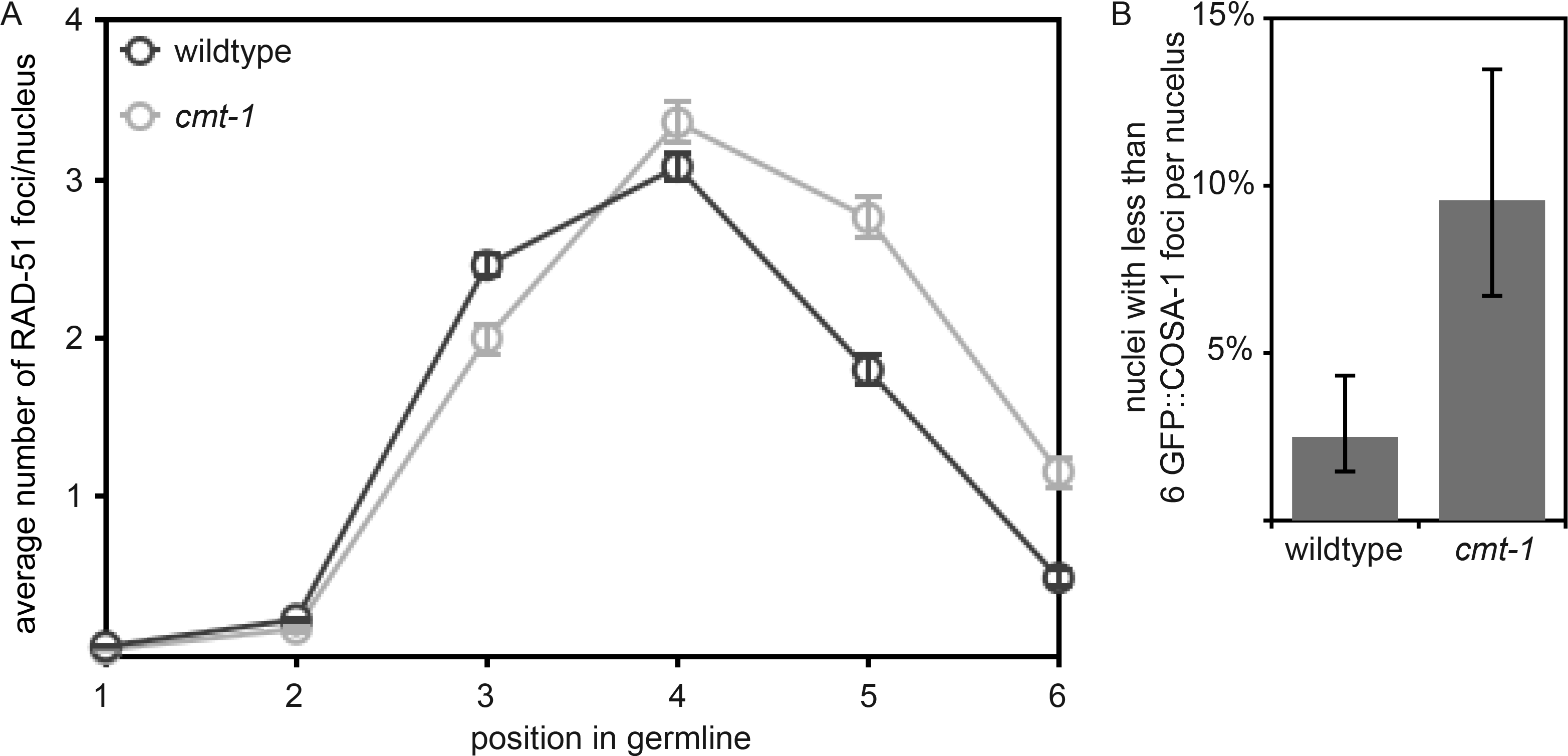
Meiotic DNA repair is not affected in *cmt-1* mutants but crossover assurance is, similar to *pch-2*^*E253Q*^ mutants. A. Timecourse of the average number of RAD-51 foci per nucleus in wildtype and *pch-2*^*E253Q*^ mutant germlines. B. Percentage of nuclei with less than six GFP∷COSA-1 foci in wildtype animals and *pch-2*^*E253Q*^ mutants.

